# LINE-1 retrotransposition is a recurrent cause of *MET* exon 14 skipping in cancer

**DOI:** 10.64898/2026.02.19.706876

**Authors:** Jennifer A. Karlow, Callan O’Connor, Radwa Sharaf, Dean C. Pavlick, Andrej Savol, Christopher Darcy, Akshay Kakumanu, William Camara, Matthew Walsh, Tyler Janovitz, Michael J. Kelley, Christine N. Serway, Jerry Mitchell, Julia A. Elvin, Meagan Montesion, Kathleen H. Burns, Garrett M. Frampton

## Abstract

*MET* exon 14 skipping is a pathogenic event that results in decreased ubiquitin-mediated degradation of the MET receptor, sustained oncogenic signaling, and conferred sensitivity to MET tyrosine kinase inhibitors. While exon 14 skipping is most commonly caused by somatically acquired base substitutions and small indels near the exon 14 splice sites, here we report nine cases in which long interspersed element-1 (LINE-1, L1)-mediated insertions within or adjacent to *MET* exon 14, including one case of a LINE-1-mediated pseudogene insertion, appear to cause exon 14 skipping. These describe the first recurrent and clinically actionable mutations caused by LINE-1 retrotransposition in cancer.

## INTRODUCTION

*MET* exon 14 skipping represents a mechanism of oncogene activation best understood in non-small cell lung cancer (NSCLC), wherein it defines a distinct and targetable subtype. These alterations are found in approximately 3% of lung adenocarcinomas and are typically mutually exclusive with mutations in *KRAS, EGFR*, and *ALK*^1–3^. Exon 14 encodes the juxtamembrane domain of the MET receptor tyrosine kinase, including a binding site for CBL E3 ubiquitin ligase, which mediates MET protein turnover^4,5^. Somatic mutations resulting in exon 14 skipping disrupt CBL binding and ubiquitination, increase MET stability, and promote sustained activation of downstream signaling pathways. *MET* exon 14 skipping is relevant to therapy selection in lung cancer, with FDA-approved MET inhibitors, such as capmatinib and tepotinib, demonstrating clinical benefit in NSCLC^2,6–8^.

Detecting DNA alterations that cause *MET* exon 14 skipping is technically challenging due to the heterogeneity of these mutations. Exon skipping can be caused by large genomic deletions that encompass the entire exon as well as a variety of base substitutions and small insertions or deletions (indels) within exon 14 or its surrounding intronic regions, which include the splice donor and acceptor sites^4,9,10^. RNA-based assays that directly capture aberrant splicing between *MET* exons 13 and 15 may improve recognition of these cases.

Long interspersed element-1 (LINE-1, L1) retrotransposition is an additional source of mutagenesis in cancers^11^, resulting in *de novo* insertions and potential retrotransposition-mediated chromosomal instability ^11–14^. Many of the most commonly occurring cancers express LINE-1 and acquire somatic LINE-1 insertions, most pervasively esophageal, lung, head and neck, and colorectal cancers^11,12,15^. LINE-1 sequences comprise 17% of the human genome and it is estimated that ∼100 individual loci are retrotransposition-competent^16^. These active LINE-1 copies are 6 kb and contain two open reading frames, ORF1 and ORF2, which encode an RNA-binding chaperone (ORF1p) and a protein with endonuclease (EN) and reverse transcriptase activities (ORF2p). These proteins form ribonucleoprotein complexes that preferentially bind the encoding LINE-1 RNA, though can associate with other RNAs in *trans*^17,18^. New insertions are initiated by ORF2p EN, which generates a genomic DNA nick that is extended by reverse transcription of the L1 RNA – a process termed target-primed reverse transcription^19,20^. Insertions are resolved with flanking target site duplications (TSDs) and are often 5′ truncated or 5′ inverted, with the 5′ segment of the LINE-1 opposing the orientation of the 3′ end^14,21^. In addition to inciting genome instability, LINE-1 insertions infrequently but recurrently drive cancers by causing loss-of-function mutations in tumor suppressor genes; 1% of colon cancers have acquired L1 insertions in an exon of *APC* ^22–24^.

## RESULTS

As part of routine clinical care, a patient with lung adenocarcinoma received combined DNA- and RNA-based next-generation sequencing (NGS) on a solid tumor biopsy. Both assays included hybrid capture of all protein-coding regions of *MET*, with the DNA assay additionally capturing the intronic regions flanking either side of exon 14. RNA results detected an in-frame fusion of *MET* exons 13 and 15, indicative of *MET* exon 14 skipping (Figure 1A, Case L1). Despite most RNA reads supporting the fusion, no causative base substitutions or indels were initially reported from DNA sequencing. However, a DNA graph-based assembly did detect a cluster of aberrant, ambiguously mapping reads within *MET* exon 14. These reads were computationally extracted and a BLASTn analysis against the NCBI nucleotide database identified them as belonging to a LINE-1 retrotransposon. Further inspection confirmed a 248 base pair (bp) 5′ truncated LINE-1 insertion 31 bp into exon 14 (Figure 1B, Case L1, Table 1, Table S1). Like many LINE-1 insertions ^11,25,26^, this included 3′ LINE-1 sequence and a poly-A tail, flanked by a short TSD (2 bp). The insertion occurred at a non-canonical site (5′-TCTT/GC-3′) as opposed to the canonical (5′-TTTT/AA-3′)^27^.

**Table 1.**
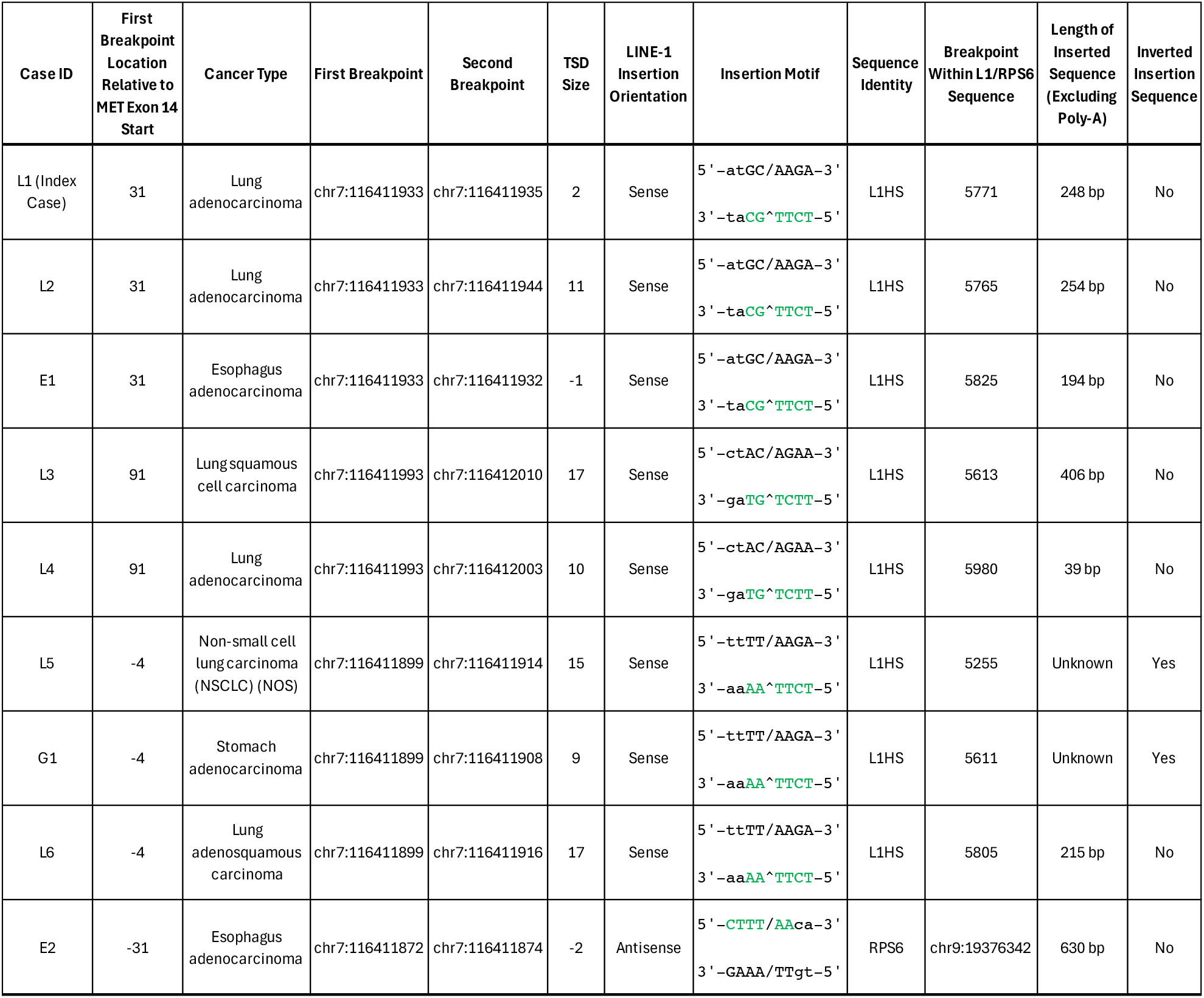
LINE-1 *MET* insertion characteristics.

**Figure 1.**
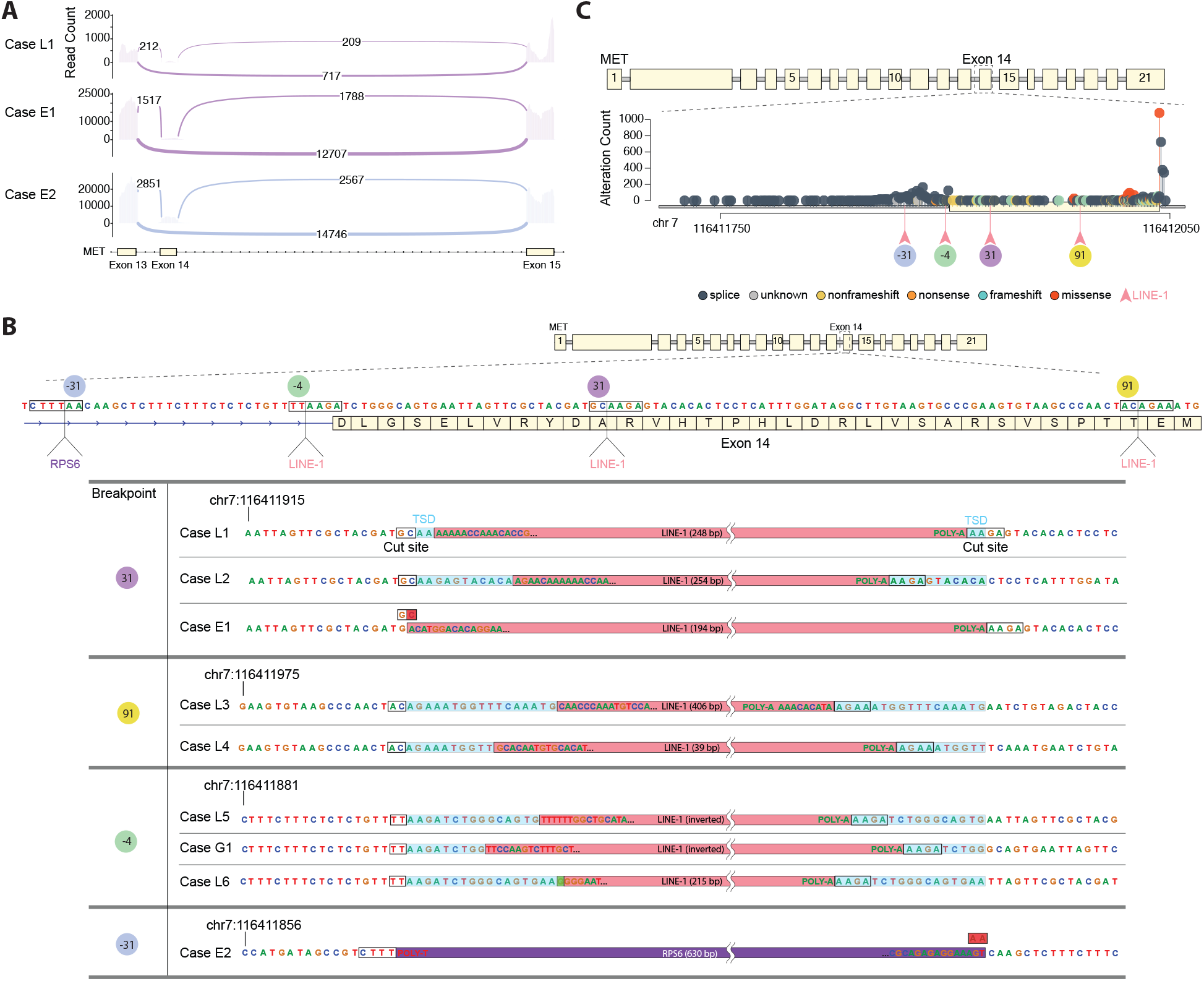
LINE-1-mediated insertions resulting in *MET* exon 14 skipping. **A**. Sashimi plots depicting RNA sequencing read coverage over *MET* exons 13, 14, and 15, as well as patterns of detected RNA splicing. Plots are shown for the index case (L1, top, purple) and two additional cases (E1, middle, purple; E2, bottom, blue). The x-axis approximates the locations of *MET* exons 13–15, represented as solid cream blocks for exons and arrows for intronic connections. The y-axis denotes read coverage at each position within an exon. Splicing junctions are illustrated as arcs connecting exons, with arc thickness being proportional to the number of supporting reads, as denoted in the arc labels. **B**. Genomic features of LINE-1-mediated insertions resulting in *MET* exon 14 skipping. Cases are organized by genomic breakpoint. Cut sites are indicated in black boxes and target site duplications (TSDs) are highlighted in blue. Red highlighted nucleotides within cut sites (shown in Cases E1 and E2) depict deleted sequence, and the green highlighted nucleotide within the Case L6 cut site depicts an inserted base pair. **C**. Lollipop plot depicting the distribution of known or likely pathogenic *MET* exon 14 variants across 2,296 tumors positive for *MET* exon 14 skipping. Each variant is colored according to alteration type: splicing (slate), unknown (gray), nonframeshift or small indels that do not disrupt the reading frame (gold), nonsense (orange), frameshift (mint), and missense (red). LINE-1-mediated insertion sites are indicated by pink arrows positioned below the axis, and their insertion location relative to the start of exon 14 is displayed below.

To investigate whether LINE-1-mediated *MET* exon 14 skipping is a recurrent phenomenon in cancer, we analyzed sequenced biopsy results from 461,707 additional patients, encompassing 96 disease groups which included 2,296 patients who were positive for previously described known or likely pathogenic *MET* exon 14 skipping alterations (Table S2). All had DNA sequencing and a subset of 11,376 cases had paired RNA sequencing. Within this cohort, we identified eight additional cases with a LINE-1-mediated insertion in or adjacent to *MET* exon 14, including five additional patients with lung cancer (Cases L2-6), two patients with esophageal cancer (Cases E1-2), and one patient with gastric cancer (Case G1) (Figure 1B, Table 1, Table S1). All insertions occurred within an interval previously characterized as a hotspot for exon skipping and all cases were devoid of other known or likely causative *MET* exon 14 skipping mutations (Figure 1C). Among all nine cases, four unique integration sites were identified: two within exon 14 and two adjacent to the splice acceptor site within intron 13.

Like the index case (Case L1), Case L2 and Case E1 exhibited a LINE-1 insertion 31 bp into exon 14 (Figure 1B, Table 1, Table S1). Case L2 had a distinct, 254 bp 5′ truncated insertion and an 11 bp TSD, supporting that this was an independently acquired insertion. Case E1 had a 194 bp 5′ truncated insertion with a 1 bp target site deletion and paired RNA sequencing that showed the dominant *MET* isoform in this biopsy contained an in-frame fusion of exons 13 and 15 (Figure 1A). Cases L3 and L4 both exhibited a LINE-1 insertion 91 bp into exon 14, at another non-canonical site (5’-TTCT/GT-3’), displaying unique inserted sequences and TSDs (Figure 1B, Table 1, Table S1). Case L3 also displayed 6 bp of transduced sequence (“CACATA”), a phenomenon observed in 20-24% of LINE-1 insertions^11,27^. This particular mobilized sequence has been previously reported and provides a fingerprint for tracing the source LINE-1 element^27^. Case L5, Case G1, and Case L6 all exhibited insertions at 4 bp upstream of the start of exon 14, at a near-canonical EN targeting motif (5′-TCTT/AA-3′) that overlaps the splice acceptor site (Figure 1B, Table 1, Table S1). Case L5 and Case G1 contained 5′ inverted LINE-1 sequences, which occur in approximately 25% of insertions^21^. The insertion in Case L6 had an untemplated guanine at the 5′ end, frequently seen in full-length insertions^28^.

The final case, E2, demonstrated an insertion at a distinct breakpoint 31 bp upstream of the start of exon 14 at another near-canonical motif (5’-CTTT/AA-3’) (Figure 1B, Table 1, Table S1). As with Cases L1 and E1, paired RNA sequencing from Case E2 showed the dominant *MET* isoform being an in-frame fusion of exons 13 and 15 (Figure 1A). However, unlike the previous cases, which all exhibited an insertion of LINE-1 DNA, the sequence in Case E2 mapped to the terminal exon of the *RPS6* gene, indicating that this was a 630 bp ORF2p-mediated pseudogene insertion. Since the DNA sequencing assay did not contain baits designed for capturing this gene, we compared the number of reads mapping to *RPS6* among all L1-positive cases. As expected, we observed an over 70-fold enrichment of *RPS6* sequences in Case E2, indicating that these reads were pulled down primarily because of its L1-mediated insertion into the baited *MET* region (Figure S1).

**Figure S1.**
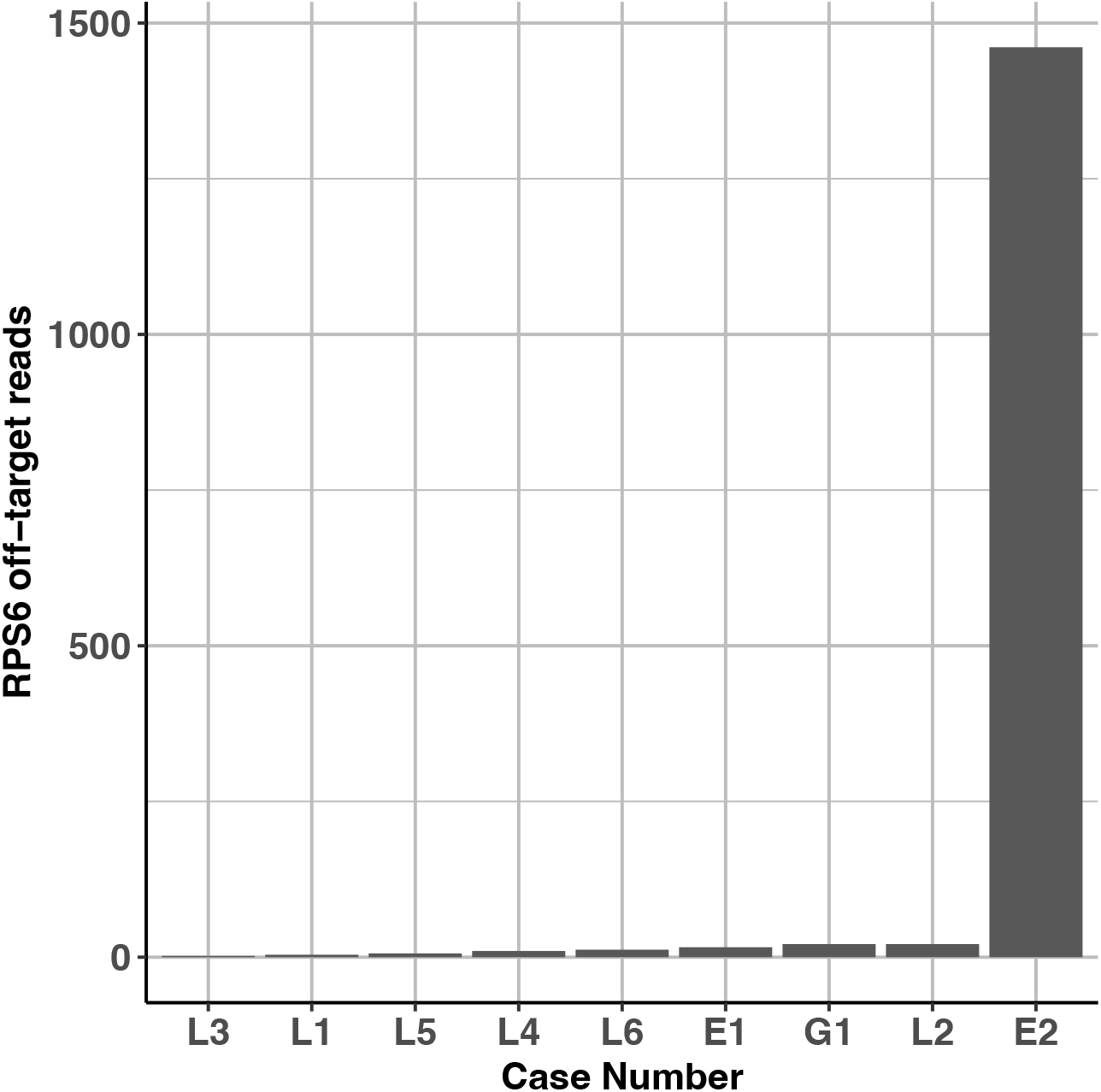
Number of reads mapping to *RPS6* per case for samples with LINE-1-mediated insertions.

## DISCUSSION

LINE-1 retrotransposition is a source of germline genetic variation in humans and an uncommon, though well-established, cause of genetic disease^29,30^. Germline LINE-1 insertions can disrupt gene function by integrating into coding exons as well as proximal to splice donor and acceptor sites, typically causing disease through loss-of-function effects.

LINE-1 can also contribute to somatic mosaicism and, in many malignancies, large numbers of somatically acquired LINE-1 insertions accumulate in cancer genomes^11^. The vast majority are not obviously deleterious to gene function — found with subclonal variant allele frequencies in non-recurrent positions in intergenic space and deep in introns — and are likely passenger mutations. Exceptions include insertions that disrupt tumor suppressor genes, most notably *APC* in colorectal cancers^22–24^. LINE-1 insertions causing oncogene activation are less common and not known to be recurrent; none previously reported are clinically actionable^31^.

In this study, we report nine instances of LINE-1-mediated insertions occurring within or adjacent to exon 14 of the *MET* oncogene, including one LINE-1-mediated pseudogene insertion, that cause exon skipping. Together, these represent a novel mutational pathway to a state of MET hyperactivation strongly implicated in driving tumorigenesis. While matched normal DNA was not available for comparison, these appear to be somatically acquired events, and all are independent insertions not known to exist as germline variants. Furthermore, genomic alterations leading to MET exon 14 skipping have never been reported in the germline.

Since *MET* exon 14 skipping is typically mutually exclusive with other oncogenic drivers in lung cancer^9,32^, we investigated co-occurring genetic alterations. Among the 9 cases, only Case L2 had a high tumor mutation burden (TMB; >10 mutations/Mb); none harbored activating mutations in *KRAS, EGFR, ERBB2*, or *BRAF*, or rearrangements in *ALK, RET, ROS1*, or *NTRK*, which is consistent with MET activation constituting a principal driver. Of note, two samples (Cases L3 and L4) harbored an *EGFR* amplification (Table S1). Additionally, we searched for *MDM2* and *CDK4* amplifications because they are reported to often co-occur with MET exon 14 skipping^9,32^. Indeed, three of the nine samples (Cases L3, L4, and L6) contained *MDM2* and/or *CDK4* amplifications.

From a clinical perspective, MET activation can be an indication for targeted therapy. Therefore, recognizing rare genetic causes of *MET* exon 14 skipping, such as LINE-1-mediated retrotransposition, may provide actionable information and underscore the utility of RNA-based testing or the development of genomic DNA assays tailored to identify these events. As clinical history was unavailable for the cases reported here, future studies are warranted to directly connect this biology to treatment response.

## METHODS

### Comprehensive Genomic Profiling

Solid tumor biopsies from 461,708 patients were collected for comprehensive genomic profiling as part of routine clinical care. Specimens were collected between January 2013 and August 2025 and processed in a Clinical Laboratory Improvement Amendments (CLIA)-certified, College of American Pathologists (CAP)-accredited, New York State-approved laboratory (Foundation Medicine, Inc.). A waiver of informed consent and Health Insurance Portability and Accountability Act waiver of authorization obtained from the Western Institutional Review Board (Protocol number: 20152817) was used for patient selection in this study cohort.

DNA-based NGS was performed on formalin-fixed paraffin embedded (FFPE) tissue sections using adapter-ligated hybridization-capture for all coding exons of up to 324 cancer-related genes, including *MET*, as previously described (FoundationOne^®^, FoundationOne^®^CDx, or FoundationOne^®^Heme (a laboratory developed test (LDT) assay)^33,34^. The specimens were sequenced to a median coverage of >500X and analyzed for short variant alterations (including base substitutions and indels), copy number alterations (gene amplifications and homozygous deletions), as well as rearrangements and gene fusions. TMB was calculated by quantifying the number of non-driver somatic coding mutations per megabase of genome sequenced^35,36^. Known or likely pathogenic variants, including those that cause *MET* exon 14 skipping, were described for each L1-positive patient (Table S1). *MET* exon 14 variant lollipop plots were generated using the trackViewer R package to visualize the spatial distribution of variants across the *MET* exon 13-15 region (chr7:116,411,709-116,412,060; transcript MET NM_000245.4, GRCh37)), with variants color-coded by functional consequence (frameshift, missense, nonsense, and splice variants).

A subset of 11,376 patients additionally received RNA-based NGS as part of routine clinical care. For these samples, RNA was co-extracted from the FFPE specimens alongside the DNA and underwent hybrid-capture-based targeted RNA sequencing using an assay designed to optimally detect gene expression and fusions, including those that cause *MET* exon 14 skipping, as previously described (FoundationOne^®^RNA, a LDT assay)^36^. Read support for *MET* exon 14 splice junctions was visualized using ggsashimi.

### Detection of LINE-1-Mediated MET Exon 14 Skipping

To identify LINE-1-mediated *MET* exon 14 skipping events from DNA-based NGS results, sequencing reads that aligned to either *MET* exon 14 or one of its adjacent introns were extracted from BAM files using samtools v1.16.1. Reads containing soft-clipped sequences greater than 5 bp in length were extracted and compiled into a FASTA file with positional annotations. The extracted reads were then aligned to the Human LINE1 (L1.3) repetitive element DNA sequence (GenBank: L19088.1) using BLASTn (NCBI BLAST+ v.2.6.0) and hits were filtered to retain only matches to the L1 sequence. The specimen with a suspected LINE-1-mediated pseudogene insertion was negative for a canonical *MET* exon 14 skipping alteration (as detected by DNA-based NGS) but positive for a fusion of MET exons 13 and 15 (as detected by RNA-based NGS). Similar to above, soft-clipped portions of reads mapping near *MET* exon 14 were extracted and subjected to a BLASTn search (taxid: 9606 (*Homo sapiens*)), revealing perfect concordance with a portion of the *RPS6* gene. All candidate L1-positive specimens were manually inspected and characterized using Integrative Genomics Viewer to identify breakpoints, TSDs, cut sites, and inserted sequences. Clipped reads at breakpoints were aligned to the consensus LINE-1 sequence (L1RP) to determine the portion of LINE-1 sequence inserted and identify any inversions.

